# Selective carbon sources influence the end-products of microbial nitrate respiration

**DOI:** 10.1101/829143

**Authors:** Hans K. Carlson, Lauren M. Lui, Morgan N. Price, Alexey E. Kazakov, Alex V. Carr, Jennifer V. Kuehl, Trenton K. Owens, Torben Nielsen, Adam P. Arkin, Adam M. Deutschbauer

**Affiliations:** Environmental Genomics and Systems Biology Division, Lawrence Berkeley National Laboratory, Berkeley, CA 94720, USA; Joint Genome Institute, Lawrence Berkeley National Laboratory, Berkeley, CA 94720, USA; Department of Bioengineering, University of California, Berkeley, CA 94720, USA; Department of Plant and Microbial Biology, University of California, Berkeley, CA 94720, USA

## Abstract

Respiratory and catabolic pathways are differentially distributed in microbial genomes. Thus, specific carbon sources may favor different respiratory processes. We profiled the influence of 94 carbon sources on the end-products of nitrate respiration in microbial enrichment cultures from diverse terrestrial environments. We found that some carbon sources consistently favor dissimilatory nitrate reduction to ammonium (DNRA/nitrate ammonification) while other carbon sources favor nitrite accumulation or denitrification. For an enrichment culture from aquatic sediment, we sequenced the genomes of the most abundant strains, matched these genomes to 16S rDNA exact sequence variants (ESVs), and used 16S rDNA amplicon sequencing to track the differential enrichment of functionally distinct ESVs on different carbon sources. We found that changes in the abundances of strains with different genetic potentials for nitrite accumulation, DNRA or denitrification were correlated with the nitrite or ammonium concentrations in the enrichment cultures recovered on different carbon sources. Specifically, we found that either L-sorbose or D-cellobiose enriched for a *Klebsiella* nitrite accumulator, other sugars enriched for an *Escherichia* nitrate ammonifier, and citrate or formate enriched for a *Pseudomonas* denitrifier and a *Sulfurospirillum* nitrate ammonifier. Our results add important nuance to the current paradigm that higher concentrations of carbon will always favor DNRA over denitrification or nitrite accumulation, and we propose that, in some cases, carbon composition can be as important as carbon concentration in determining nitrate respiratory end-products. Furthermore, our approach can be extended to other environments and metabolisms to characterize how selective parameters influence microbial community composition, gene content and function.

## Introduction

The nitrogen cycle is the most anthropogenically perturbed element cycle(*1*). The world’s population relies on nitrogen fertilizer to maintain productive agricultural ecosystems. However, nitrogen contamination of surface waters and groundwater has serious consequences for public and environmental health. Thus, we need a predictive understanding of how varying biogeochemical conditions influence the fate of nitrogen in the environment. Such a framework would enable better monitoring and managing of microbial communities to mitigate environmental damage while maximizing the productivity of agricultural lands(*2*).

Heterotrophic nitrate respiration is a critical juncture in the nitrogen and carbon cycles. Denitrification converts nitrate into dinitrogen, thereby returning biologically or industrially fixed nitrogen to the atmosphere. Dissimilatory nitrate reduction to ammonium (DNRA, nitrate ammonification) converts nitrate into ammonium, thereby maintaining nitrogen in terrestrial reservoirs. Most studies agree about the importance of electron donor excess or limitation in controlling the competition between DNRA and denitrification(*3*-*11*). There are some examples of different electron donors (e.g. carbon sources) determining the end-products of nitrate respiration(*5, 8*-*10*), but less is known about the mechanistic basis for these observations. While thermodynamic calculations predict that higher concentrations of electron donor will favor DNRA over denitrification(*7*), the specific carbon source available to drive DNRA must be utilized by the DNRA sub-populations in the system. Nitrate respiratory pathway enzymes are differentially distributed across phylogenetic boundaries(*12*) as are carbon catabolic pathways (*13, 14*). Thus, we postulate that certain, selective carbon sources are more likely to drive microbial nitrate respiration towards specific end-products such as dinitrogen (N_2_), ammonium (NH_4_^+^) or intermediate nitrogen oxides (NO_2_^−^, NO, N_2_O), especially in systems with less complex microbial communities.

Carbon amendments are often used to perturb microbiomes to alter community composition and functional outcomes(*9, 15*-*20*). Usually, however, carbon sources are chosen based on general physiological hypotheses (e.g. acetate as a non-fermentable carbon source to stimulate metal reduction(*18*), poly-L-lactate as a hydrogen releasing compound to stimulate reductive dehalogenation(*20*)), and rarely are carbon sources systematically compared to identify the optimal carbon source to favor a given function. Some studies suggest that different carbon sources will enrich for microbial sub-populations with distinct carbon catabolic preferences (*21*-*24*), but the mechanisms and functional consequences of these changes in microbiome composition on key ecosystem services remain largely uncharacterized. While advances in high-throughput genetics are leading to more rapid discovery of genes involved in carbon catabolic pathways(*25*), our current ability to predict carbon preferences based on taxonomy remains poor, particularly when catabolic preferences vary for closely related taxa(*13, 14*). Additionally—even with data on the genetic potential of microbial sub-populations from genome sequencing—inaccurate gene annotations, complex gene regulation, ecological dynamics and environment-specific physiological and metabolic responses make taxonomy-based or genome-based predictions of community composition and functional traits difficult. To bridge this knowledge gap, there is a need for high-throughput methods to rapidly measure the influence of selective pressures on microbial community composition, gene content and function across diverse conditions in the laboratory(*26*).

High-throughput colorimetric assays to measure microbial activity can be combined with 16S rDNA amplicon sequencing of enrichment cultures to understand how changes in community composition influences metabolic traits (Figure 1A)(*26*-*29*). By determining the gene content of each strain represented by a 16S rDNA exact sequence variant (ESV) in a microbial community using genome-resolved metagenomics and isolate genome sequencing, we can track changes in gene content in high-throughput using 16S rDNA amplicons (Figure 1B). Thus, we can measure correlations between growth conditions, strain abundances, functional gene abundances, and functional traits (Figure 1) to understand how selective growth conditions influence the functional ecology of a microbial community.

**Figure 1.**
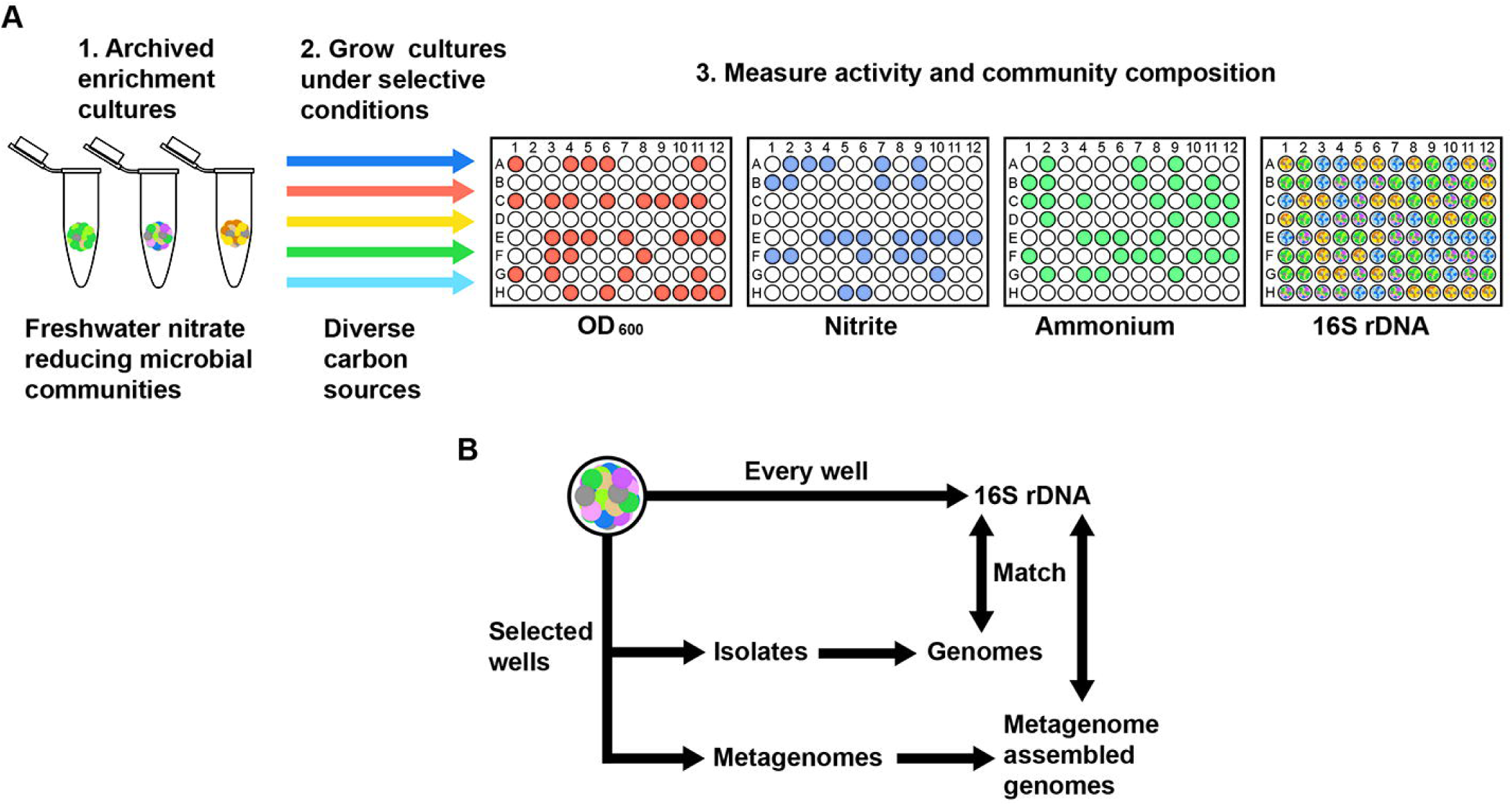
Workflow to measure the influence of selective growth conditions on microbial community composition, gene content and functional activity. **A.** Archived microbial enrichment cultures are cultured under different growth conditions. Community functional traits, community composition and both strain and community genetic potential are measured. In the present work, freshwater nitrate reducing microbial communities are grown on 94 different carbon sources, some of which are selective for different end-products or intermediates of nitrate reduction. Growth (optical density/OD 600), nitrite and ammonium are measured through colorimetric assays, and microbial community composition is determined using 16S rDNA amplicon sequencing. **B.** Pure culture microbial isolates, isolate genomes and metagenome assembled genomes (MAGs) are obtained for select cultures. Matches between 16S rDNA sequences in genomes and MAGS with amplicons allows the assignment of genetic potential to the 16S rDNA exact sequence variants (ESVs) in all of the enrichment cultures.

In this study, we characterized the influence of 94 different carbon sources on nitrate-respiring microbial communities. We used colorimetric assays to quantify nitrite and ammonium concentrations, and we identified carbon sources that favor different end-products of nitrate respiration across microbial communities from diverse environments. We then focused on a microbial community enriched from aquatic sediment. We recovered this enrichment culture on different carbon sources and observed correlations between high nitrite or ammonium concentrations and high relative abundance of specific strains with the genetic potential to produce these end-products. We found that D-glucose favors ammonium production and the growth of an *Escherichia* community member with the genetic potential for DNRA, but L-sorbose favors nitrite accumulation and selects for a *Klebsiella* nitrite accumulator. In contrast, citrate or formate enrich for a *Pseudomonas* denitrifier and a *Sulfurospirillum* nitrate ammonifier. Isolation and characterization of strains from the enrichments confirms the catabolic and respiratory traits predicted from sequencing the genomes of strains in the community. Finally, comparative genomic analyses suggest that our findings with L-sorbose may be a likely outcome across other environments. Taken together, our results indicate that alongside carbon concentration, carbon composition influences the end-products of nitrate respiration by enriching for sub-populations with distinct respiratory traits. This approach to linking selective carbon sources to changes in the composition and gene content of a nitrate-respiring microbial community can be extended to other systems and microbiomes to characterize of how carbon sources and other selective pressures influence the functional ecology of this, and other, globally important metabolic processes.

## Materials and Methods

### Media and cultivation conditions

Samples for primary enrichments were sediment collected from Jewel Lake in Tilden Regional Park (37°54’45.2”N 122°16’09.1”W), soil from the Russell Ranch Field Site (38°32’38.8”N 121°52’12.4”W), or groundwater from the Oak Ridge Field Research Center (35°56’27.8484”N 84°20’10.2516”W). Primary microbial enrichment cultures were prepared by mixing sediment or soil (∼10 grams) or groundwater (5 mL) with anoxic chemically defined basal medium supplemented with 2 g/L yeast extract (Becton Dickinson and Company, Franklin Lakes, NJ, USA) as the sole organic carbon source and electron donor and 20 mM sodium nitrate as the sole terminal electron acceptor and incubated for 48 hours at 30 °C. All chemicals are from Sigma-Aldrich (St Louis, Mo, USA). Basal medium contained per liter: 1 g sodium chloride, 0.25 g ammonium chloride (4.67 mM), 1 g sodium phosphate, 0.1 g potassium chloride and 30 mM HEPES buffer with vitamins and minerals added from 100x stock solutions. Vitamin stock solution contained per liter: 10 mg pyridoxine HCl, 5 mg 4-aminobenzoic acid, 5 mg lipoic acid, 5 mg nicotinic acid, 5 mg riboflavin, 5 mg thiamine HCl, 5 mg calcium D,L-pantothenate, 2 mg biotin, 2 mg folic acid, 0.1 mg cyanocobalamin. Mineral stock solution contained per liter: 3 g magnesium sulfate heptahydrate, 1.5 g nitrilotriacetic acid, 1 g sodium chloride, 0.5291 g manganese(II) chloride tetrahydrate, 0.05458 g cobalt chloride, 0.1 g zinc sulfate heptahydrate, 0.1 g calcium chloride dihydrate, 0.07153 g iron(II) chloride tetrahydrate, 0.02765 g nickel(II) sulfate hexahydrate, 0.02 g aluminum potassium sulfate dodecahydrate, 0.00683 g copper(II) chloride dihydrate, 0.01 g boric acid, 0.01 g sodium molybdate dihydrate, 0.000197 g sodium selenite pentahydrate. Enrichments were passaged twice by ten-fold dilution into fresh basal medium and cryopreserved in multiple aliquots in basal medium with nitrate but without yeast extract and containing 25% glycerol.

To measure the influence of carbon sources on the end-products of the archived nitrate reducing microbial communities, cryo-preserved enrichments were recovered in anoxic chemically defined basal medium amended with 2g/L yeast extract and 20 mM sodium nitrate. Cells from recovered enrichment cultures were pelleted at 4000 RCF and washed three times with 2x concentrated basal medium lacking a carbon source. Washed cells were resuspended in 2x concentrated basal medium lacking a carbon source to an optical density (OD 600) of 0.04 and the cell suspension was transferred into either 384 well microplates (Costar, Thermo Fisher Scientific, Waltham, MA, USA) or 96 deep-well blocks (Costar) in which 94 carbon sources and water controls were arrayed (Table S1). Carbon source stock solutions were added to microplates using a Biomek FxP liquid handling robot (Beckman Coulter, Indianapolis, IN, USA) and kept in an anaerobic chamber (Coy, Grass Lake, MI, USA) for 48 hours to become anoxic prior to inoculation using a Rainin Liquidator 96 pipettor (Mettler-Toledo, Oakland, CA, USA). Inoculated microplates were sealed with silicon microplate seals (VWR) and incubated at 30 °C in an incubator in an anaerobic chamber (Coy). Growth was monitored by optical density (OD 600) using a Tecan M1000 Pro microplate reader (Tecan Group Ltd., Männendorf, Switzerland) and cultures were harvested at 48 hours for DNA sequencing and colorimetric assays to measure nitrogen cycle metabolic intermediates.

### Isolation of bacterial strains

To obtain pure culture isolates, liquid enrichments were recovered in anoxic basal medium containing 20 mM sodium nitrate and amended with carbon sources in which target strains were enriched. Liquid cultures were then plated onto anoxic solid agar containing the same media. Colonies were picked into either basal medium or R2A medium and recovered either aerobically or anaerobically with 20mM sodium nitrate as the sole terminal electron acceptor. Isolates were cryo-preserved in 25% glycerol and DNA was extracted for genome sequencing and 16S rDNA Sanger sequencing.

### Colorimetric assays and analysis

Nitrite and ammonium concentrations were determined using established colorimetric assays(*30*). Microplate seals were removed from 384-well microplates containing enrichment cultures and a Biomek FxP (Beckman Coulter) was used to transfer small volumes of culture to assay microplates pre-filled with small volumes of ultrapure water. For nitrite measurements, 2 *µ*L of culture in 20 *µ*L of water was prepared in assay plates. 20 *µ*L Griess reagent was added to assay plates which were then kept at 30 °C for 30 minutes prior to reading absorbance at 548 nm (Tecan M1000 Pro). Greiss reagent contains 0.2% w/v napthylethylenediamine dihydrochloride, 2% w/v sulfanilamide and 5% phosphoric acid. For ammonium measurements, 4 *µ*L of culture diluted in 20 *µ*L of distilled deionized water was prepared in assay plates. In sequential order, 4 *µ*L of citrate reagent, 8 *µ*L of salicylate/nitroprusside reagent and 4 *µ*L bleach reagent were added to assay plates which were then kept at 30 °C for 30 minutes. Citrate reagent contains 10 g trisodium citrate and 4 g sodium hydroxide in 200 mL water. Salicylate/nitroprusside reagent contains 15.626 g sodium salicylate and 0.250 g sodium nitroprusside in 200 mL water at pH 6-7. Bleach reagent contains 1g sodium phosphate monobasic, 2 mL 2M sodium hydroxide, 10 mL bleach (0.7 M NaOCl, Chlorox Company, Pleasanton, CA, USA) in 100 mL water at pH 12-13. All reagents were prepared the same day as assays and standard curves with sodium nitrite and ammonium chloride were used to calculate nitrite and ammonium concentrations. For pH measurements, 100 *µ*M resazurin was mixed 1:1 with cultures and absorbance was measured at 590 nM. A standard curve was prepared in sterile media with different buffer salts to cover the pH range from 3 to 11 as reported previously*(31*). For all colorimetric assays, we also confirmed that interference of all 94 carbon sources was negligible. Using constants obtained from the BioNumbers database(*32*), we estimated the quantity of nitrogen assimilated into biomass by assuming 0.3 g/L of dry weight of bacterial culture at OD 600 = 1 (BNID 109835, BNID 109836) (*33, 34*), and by assuming 12% nitrogen by weight in microbial biomass based on measured C:N:P ratios(*35, 36*).

### 16S rDNA amplicon sequencing and analysis

DNA extraction, library prep and Illumina sequencing were performed as reported previously(*37*). Briefly, microbial cells from 500 *µ*L cultures were pelleted by centrifugation at 4000 RCF after 48 hours of growth at 30°C. Genomic DNA extractions were performed using the QIAamp 96 DNA QIAcube HT Kit (Qiagen, Redwood City, CA, USA) with minor modifications including an enzymatic lysis pre-treatment step and the use of a vacuum manifold to perform column purification steps.

Following gDNA extraction, gDNA concentrations were quantified using the Quant-iT dsDNA High-Sensitivity kit (Thermo Fisher Scientific, Waltham, MA, USA) and normalized to approximately 3 ng/*µ*l. PCR amplification of the V3–V4 region of the 16S rDNA gene was performed with Phusion High-Fidelity DNA Polymerase (New England Biolabs, Ipswich, MA, USA) for 25 cycles using 0.05 M of each primer as described previously(*37*). PCR amplicons were pooled by plate (96 conditions), purified (Zymo Research, Irvine, CA, USA), and quantified using the Quant-iT dsDNA High-Sensitivity kit (Thermo Fisher). The samples were normalized to the lowest sample concentration and then combined in equal proportions to generate the library. The library was quantified prior to loading using quantitative real-time PCR (KAPA Biosystems, Wilmington, MA, USA) on a CFX96 real-time PCR detection system (Bio-Rad, Hercules, CA, USA). Following amplification, the library was diluted to 4.5 nM and loaded on the Illumina MiSeq platform for 2 × 300 bp paired-end sequencing.

To obtain exact sequence variants (ESVs) from the 16S amplicon sequencing data, we used QIIME2 v2018.2. Primers were trimmed from Illumina reads using custom scripts prior to QIIME2 processing. Reads were discarded if primers were not detected or did not have a matching paired read. The DADA2 pipeline was used to identify ESVs and to create a relative abundance table with 280 for the --p-trunc-len-f and --p-trunc-len-r parameters for read quality trimming. We focused on ESVs that were present at >5% in any sample. The fold-enrichment of each strain on each carbon source relative to the primary enrichment inoculum is reported in Table S3. *Sulfurospirillum* was below detection in the inoculum and we calculated a lower limit for the fold enrichment of this strain based on the observation that the lowest abundance ESVs in our samples were observed at ∼0.01%.

### Genome and Metagenome sequencing and analysis

For isolates, we prepared sequencing libraries using the KAPA HyperPrep kit (Roche, Basal, Switzerland) and sequenced on the Illumina HiSeq2500 (Illumina, San Diego, CA, USA). Genomes were assembled using Unicycler(*38*). Cultures used for metagenome sequencing were grown on citrate, formate, L-arginine, pyruvate, lactate and yeast extract, sequenced separately and reads were combined for a co-assembly. gDNA was prepped for metagenomics sequencing using the Nexterra Flex kit (Illumina) and sequenced on the Illumina HiSeq2500 with 2×150 paired-end reads. Metagenome reads were assembled using SPAdes v3.13.0(*39*). Protein-coding genes were predicted using PRODIGAL and RNA genes using INFERNAL v1.1. Assembled contigs were binned using MetaBat2(*40*). 16S rDNA exact sequence variants (ESVs) from amplicon sequencing of enrichment cultures were searched against microbial isolate genomes to identify exact matches. The taxonomy of the metagenome assembled genomes (MAGs) for *Sulfurospirillum, Clostridium* and *Peptostreptococcae* was determined based on the GTDB-Tk(*41*) taxonomy and matched to the 16S rDNA ESVs with the closest taxonomy from SILVA(*42*) and highest coverage. All bins used for functional assignments of strains were >69% complete as assessed by CheckM(*43*) (Table S4).

Genes involved in nitrogen cycling were identified by comparison with a manually curated database of marker proteins for nitrogen cycle processes. To construct the database, nitrogen cycle-related genes were collected from SEED (“Denitrification”, “Dissimilatory nitrite reductase”, “Nitrate and nitrite ammonification” subsystems) (*44*) and KEGG ORTHOLOGY (M00175, M00528, M00529, M00530, M00531, M00804 modules) (*45*) databases. Additional nitrogen cycle enzymes were identified by CD-HIT (*46*) clustering of the annotated nitrogen cycle enzymes with proteins from 11384 genomes from SEED database at 80% sequence identity threshold.

MAG genes related to nitrogen cycle enzymes were identified by DIAMOND(*47*) search (e-value threshold 10^−5^, minimum 50% identity) against the marker proteins database. To remove spurious homologs, all candidate genes were used in a second DIAMOND search (e-value threshold 10^−4^) against proteins from 11384 SEED genomes, not related to nitrogen cycle genes. Genes having higher bit-score in the second search were discarded as false-positives.

### Co-occurrence of L-sorbose utilization genes with nitrate reduction genes

We used Annotree(*48*) to search 28,941 prokaryotic genomes from GTDB-Tk(*41*) for KEGG orthologs of genes involved in L-sorbose utilization(*49*) (*sorABE:* KO2814, KO2813, K19956), respiratory nitrate reduction (*narG:* KO00370, *napA*: KO2567) and respiratory nitrite reduction (*nirS*: K15864, *nirK*: KO0368, *nrfA:* KO3385). We used default settings on Annotree (minimum 30% amino acid identity between subject and a gene assigned to that ortholog group by KEGG).

## Results

### Selective carbon sources influence the end-products of nitrate respiration in microbial enrichment cultures

We enriched for nitrate-respiring microbial communities by inoculating anoxic basal growth medium with 20 mM sodium nitrate as the sole terminal electron acceptor and 2g/L yeast extract as the sole carbon source and electron donor with aquatic sediment from Jewel Lake in Tilden Regional Park in Berkeley, CA, agricultural soil from Russell Ranch in Davis, CA and groundwater from the Oak Ridge Field Research Center in Oak Ridge, TN (ORFRC). We minimized passaging of the enrichments in an effort to preserve as diverse a community as possible. Each enrichment was cryopreserved, recovered in media with yeast extract, washed, and subsequently cultured in the presence of 94 different carbon sources. Growth, pH, nitrite and ammonium concentrations were measured after 48 hours. Because our growth medium contains ammonium as a nitrogen source, we corrected for ammonium assimilated into biomass using conversion factors based on assumptions about the percentage of nitrogen in biomass and measurements of optical density (Materials and Methods). This correction has only a minor impact on the relative ranking of carbon sources in terms of ammonium production, but for some carbon sources our estimates suggest that more ammonium was consumed than produced by the microbial community (Figure 2).

**Figure 2.**
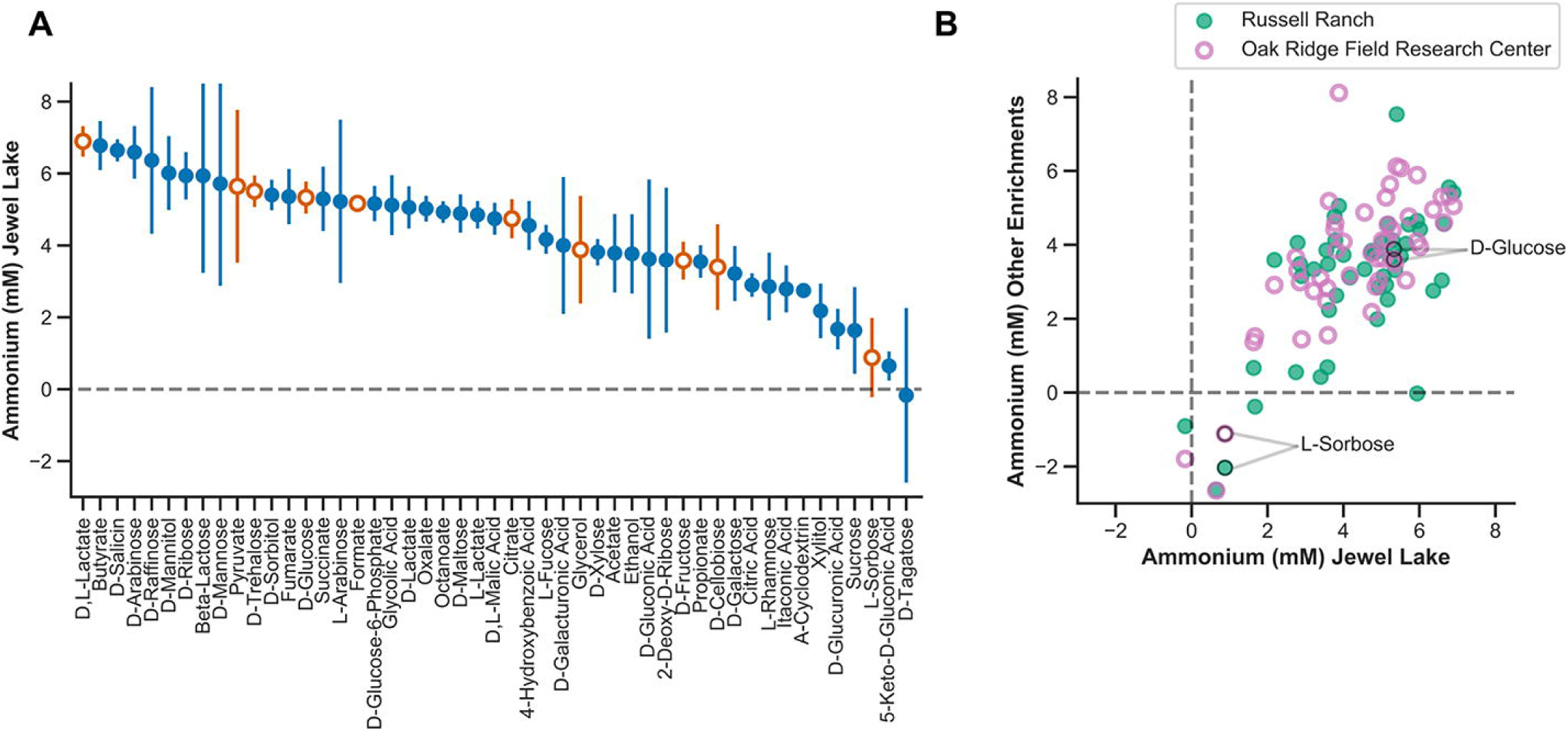
Influence of selective carbon sources on ammonium production in enrichment cultures. **A.** Ammonium production (mM) of the Jewel Lake enrichment cultured on 48 diverse sugars, organic acids and alcohols. Symbols represent means and error bars represent the standard deviation of four replicates. Orange, open symbols are carbon sources selected for further 16S microbial community analysis, isolations and metagenomics. **B.** Mean ammonium produced (mM) by cultures cultured on the 48 diverse carbon sources (shown in panel A) compared between the Jewel Lake enrichment and the Russell Ranch (closed symbols) or Oak Ridge Field Research Center enrichments (open symbols).

We were primarily interested in identifying carbon sources that influence the end-products of nitrate respiration because they are specifically utilized by microbial sub-populations with different respiratory pathways. As such, we were concerned that (1) ammonium might be released from some nitrogen containing carbon sources, (2) low pH toxicity might select against some strains, or (3) optical density measurements used to estimate ammonium assimilated into biomass might be skewed by compound precipitation. Thus, we excluded from further analysis those carbon sources that (1) contain a nitrogen atom that can be released through microbial catabolism, (2) lead to a pH < 5 after 48 hours of growth, or (3) resulted in a measurable optical density in the absence of microbial growth. In general, the carbon sources we excluded based on these criteria produced a similar range of ammonium concentrations as those we pursued in more depth (Table S1), but we expect them to have indirect effects on ammonium production and community composition aside from selecting for strains with distinct carbon catabolic and nitrate respiratory pathways. For example, in cultures amended with some amino acids and nucleotides, ammonium production was higher than was possible via reduction of the 20 mM nitrate in our growth medium alone. This is likely because ammonium is released via catabolic deamination of these nitrogen-containing carbon sources. Thus, to avoid this complicating activity, we focused on a subset of 48 carbon sources including organic acids, alcohols and sugars for further analysis (Figure 2).

We compared the ammonium concentrations in cultures grown on different carbon sources and identified carbon sources with a consistent influence on ammonium production within a single enrichment (Figure 2A) and between enrichments (Figure 2B). For example, L-sorbose reproducibly drives less ammonium production compared to D-glucose across replicates in the Jewel Lake (JL) enrichment (Figure 2A). A similar result was observed for the other two enrichments (Figure 2B, Supplemental Table S1).

There is no obvious relationship between the chemical class of carbon source and ammonium production, as both organic acids and sugars are distributed across the range of ammonium concentrations we observed (Figure 2A). Because all carbon sources were added at a concentration of 20 mM, this represents an electron equivalent excess of donor relative to acceptor (20 mM nitrate) in nearly every case regardless of whether we consider the 5 electron reduction of nitrate to dinitrogen or 8 electron reduction of nitrate to ammonium. Thus, we expect that these growth conditions should tend to favor DNRA^3-7^. As such, the clear difference in ammonium production between carbon sources with similar electron donor equivalencies demonstrates a selective influence of the carbon source on nitrate reduction end-products rather than an influence of carbon to nitrate ratio. For example, both D-glucose and L-sorbose provide 24 electron equivalents per mole, but D-glucose consistently led to more ammonium production (Figure 2).

### Microbial community compositional shifts associated with different end-products of nitrate respiration

We hypothesized that the difference in ammonium production between enrichment cultures recovered on different carbon sources could be attributed to differences in the composition and gene content of the nitrate-respiring microbial communities selectively enriched on each carbon source. To understand the relationship between ammonium production and microbial community composition, we cultured the JL enrichment in triplicate on 10 carbon sources that produced varying levels of ammonium (open symbols in Figure 2A) and measured pH, optical density, nitrite, ammonium and microbial community composition by 16S rDNA amplicon sequencing.

We measured correlations between ammonium, nitrite, optical density and pH across the enrichment cultures (Figure 3A,3B). Nitrite and ammonium concentrations are negatively correlated with each other across cultures (Pearson’s correlation r =−0.58, p = 0.00072) (Figure 3A, 3B). Also, higher nitrite concentrations are associated with lower growth (Figure 3B). This is consistent with the fact that nitrate reduction to nitrite yields less energy than nitrate reduction to ammonium or dinitrogen.

**Figure 3.**
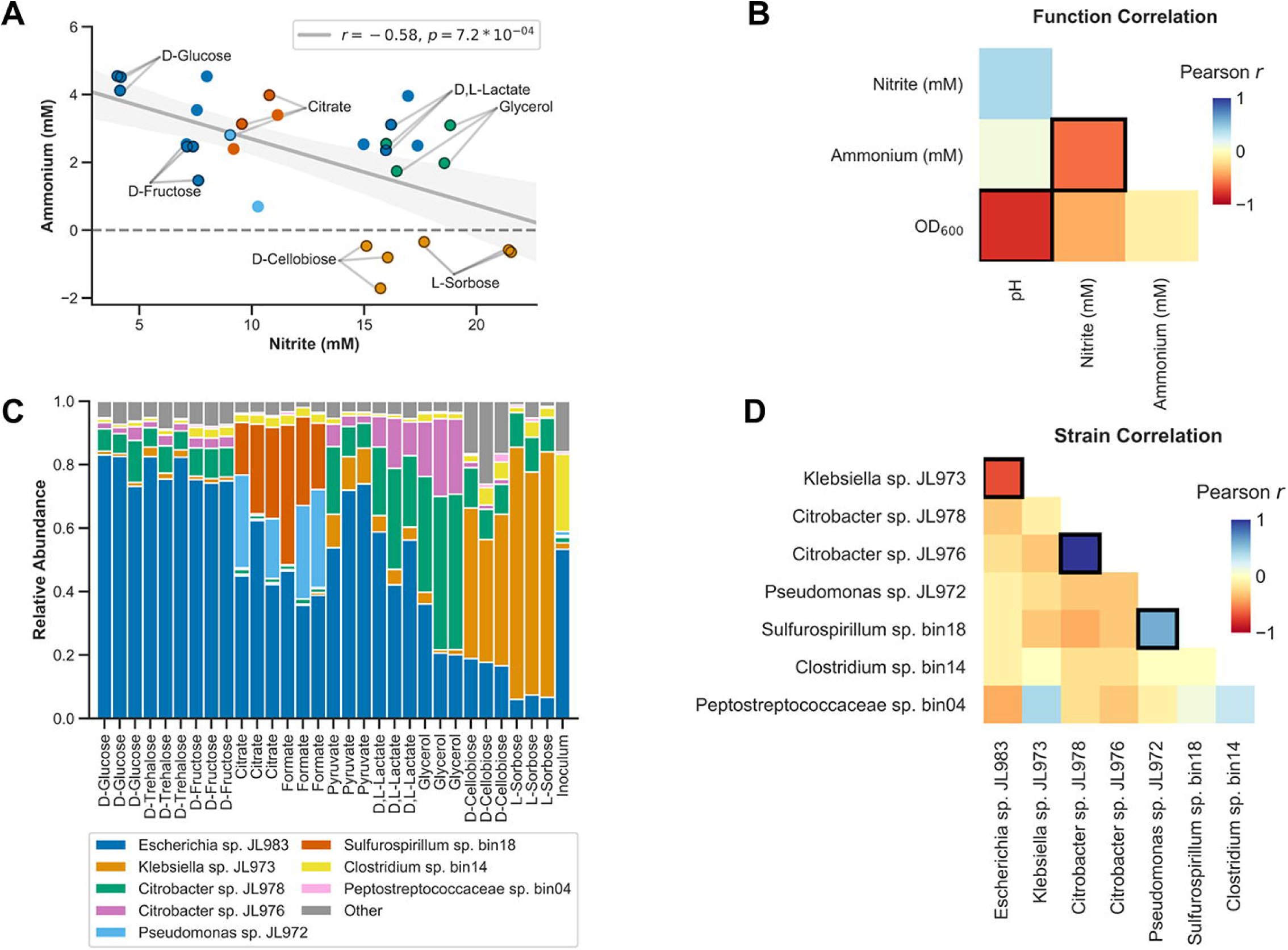
Correlations between activity measurements and between strain abundances in the Jewel Lake enrichment. **A.** Ammonium and nitrite concentrations for the Jewel Lake enrichment cultured on 10 different carbon sources in triplicate. Points are colored based on which dominant strain is most highly selectively enriched in each condition (see panel C legend). Dominant strains are 16S rDNA exact sequence variants (ESVs/strains) observed at a relative abundance of greater than 5% in any culture. Linear fit (Pearson correlation) of the nitrite and ammonium data is displayed (grey line) as well as the estimated 95% confidence interval (light grey shading) and linear correlation using Pearson’s *r* (legend). **B.** Pearson correlations between functional activity measurements for the Jewel Lake enrichment cultured on the same 10 carbon sources from panel A. Significant correlations, where *p* < 0.05 and Benjamini–Hochberg false discovery rate (FDR) *q* values were <0.1, are indicated by bold borders. **C.** Relative abundances of strains in the Jewel Lake enrichment cultured on different carbon sources. Coloring as in Panel A. **D.** Pearson correlations between the relative abundances of strains in the Jewel Lake enrichment. Significant correlations, after FDR correction, are indicated by bold borders.

The electron donor equivalents per mole provided by this set of 10 carbon sources varies from 6 electrons for formate to 44 electrons for trehalose. Thus, in most cases there is an electron equivalent excess of carbon relative to nitrate (NO_3_^−^ to N_2_ is 5 electrons, NO_3_^−^ to NH_4_^+^ is 8 electrons). We observed poor correlations between electron donor equivalents and ammonium or nitrite concentrations (Figure S1A, S1B). We also observed poor correlations between pH and ammonium or nitrite concentrations (Figure S1C, S1D). However, lower pH is associated with more growth which is consistent with organic acid production through fermentation of the sugars (Figure 3B). It is known that fermentation can compete with nitrate respiration to influence nitrate respiratory end-products(*5, 50*), but in our enrichments this is not a dominant factor.

To understand the mechanistic basis for these correlations we obtained pure cultures and sequenced the genomes of several dominant 16S rDNA exact sequence variant (ESVs) by plating the JL enrichment on anaerobic agar plates amended with carbon sources and nitrate. We also sequenced metagenomes of carbon-source enrichments that were dominated by ESVs that we did not isolate. We matched each 16S rDNA exact sequence variant (ESV) with 16S rDNA sequences in genome sequenced isolates. For ESVs we did not isolate, we matched the SILVA(*42*) taxonomy of ESVs with the GTDB-Tk(*41*) taxonomy of the most closely related metagenome assembled genome (MAG) with the highest fold coverage. For these metagenomes from low complexity enrichments, there is no ambiguity about which MAG corresponds to which 16S ESV. Thus, we are able to track the abundance of specific strains in the JL enrichment with known genetic potential across cultures using 16S rDNA amplicons.

We observed specific strains enriched on different carbon sources (Figure 3C, Figures S2). This is consistent with our hypothesis that selective carbon sources alter the composition of the microbial community and thereby influence nitrite and ammonium production. We focused on strains that are present at >5% relative abundance in any of the enrichment cultures and measured correlations between these strains (Figure 3D). The *Escherichia* and *Klebsiella* strains are strongly negatively correlated with each other (Figure 3D). The *Escherichia* strain is dominant in D-glucose, D-fructose and D-trehalose while the *Klebsiella* is dominant in L-sorbose and D-cellobiose (Figure 3C). The two *Citrobacter* strains are co-enriched on D,L-lactate and glycerol, while *Pseudomonas* and *Sulfurospirillum* are co-enriched on citrate and formate. *Clostridium* and *Peptostreptococcaceae* are below 5% relative abundance in most samples, but are more abundant in the primary yeast extract enrichment, likely because they are specialists in peptide and amino acid catabolism (Figure 3C, Table S3).

### Correlations between microbial community genetic functional potential and functional activity

We identified correlations between the relative abundances of the dominant strains and pH, OD 600, nitrite or ammonium (Figure 4A). From the metagenomic and isolate genome sequencing we know the genetic potential of all dominant strains (Figure 4B, Table S2-S4). In most cases, the strains that are positively correlated with ammonium production or nitrite production have the genetic potential to carry out that function (Figure 4A, Figure 4B Table S2-S3). For example, the *Escherichia* strain, whose abundance is positively correlated with ammonium production across our enrichments (Pearson correlation, r =0.77, p<0.0001), has the complete pathway for DNRA. In contrast, the *Klebsiella* strain, which is positively correlated with nitrite accumulation (Figure 4A), has a NarG-type nitrate reductase, but no downstream enzymes involved in DNRA or denitrification and is thus predicted to be a nitrite accumulator (Figure 4B). The *Sulfurospirillum, Pseudomonas* and *Citrobacter* strains are weakly positively correlated with ammonium production, and, with the exception of the *Pseudomonas*, all have the capacity for nitrate ammonification. The only strain with a complete denitrification pathway is the *Pseudomonas*, but the *Sulfurospirillum* has a nitrous oxide reductase (*nosZ*) and thus may participate in the final step of denitrification as well as DNRA.

**Figure 4.**
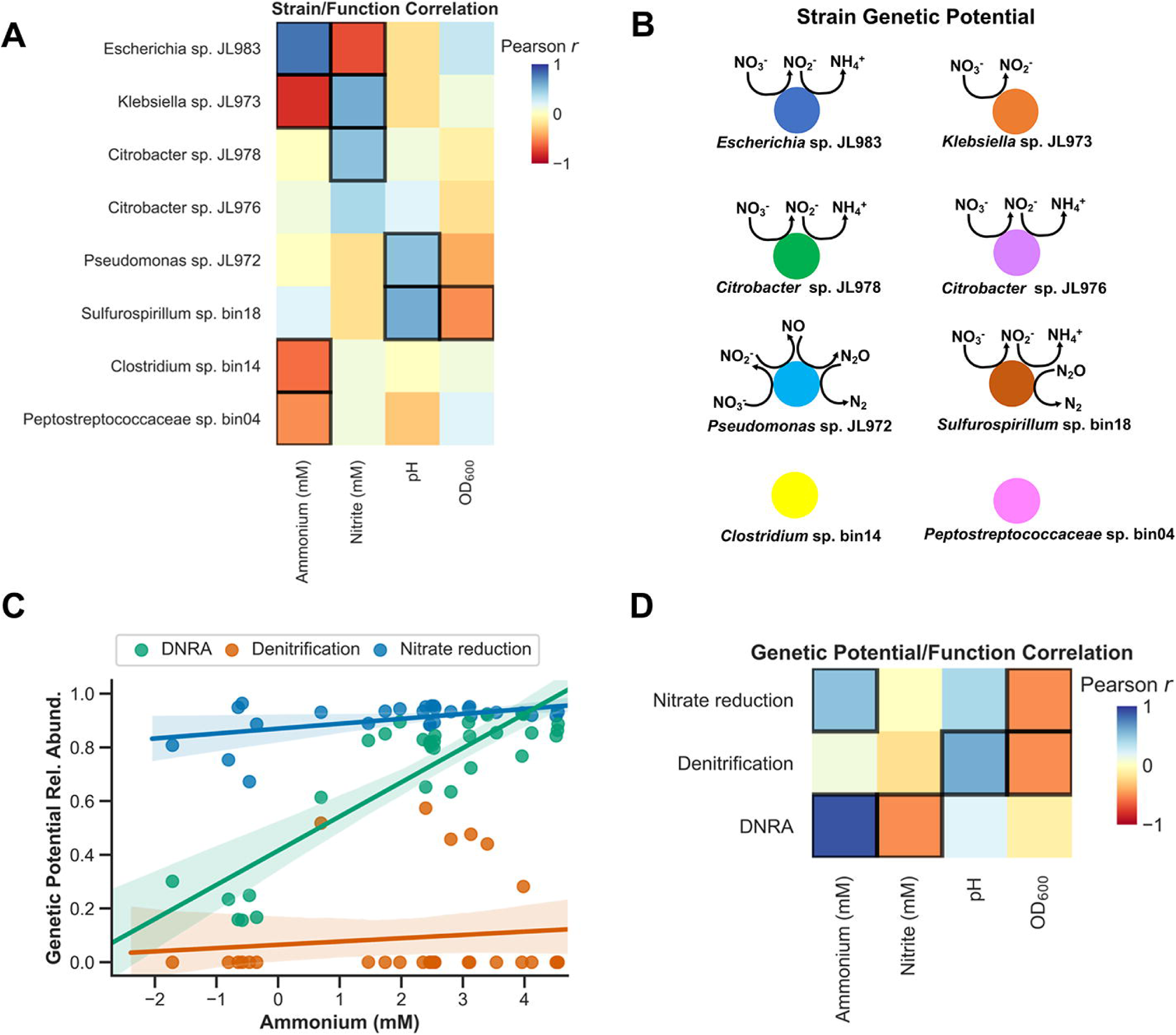
Correlations between strain abundance or total genetic potential with activity measurements in the Jewel Lake enrichment. **A.** Pearson correlations between strain relative abundances and measurements of nitrite, ammonium, pH, and OD 600. Significant correlations, after FDR correction, are indicated by bold borders. **B.** The predicted genetic potential of each strain in the Jewel Lake enrichment to catalyze steps in nitrate reduction, DNRA and denitrification. Coloring as in Figure 3C. **C.** Relative abundances of the total genetic potential for nitrate reduction, DNRA, or denitrification reduction plotted against ammonium concentrations. Each point represents a different culture. Genetic potential for each trait is the presence of genes essential for each trait. Total genetic potential is the sum of the relative abundances of each strain with each trait. **D.** Pearson correlations between the relative abundances of the genetic potential for DNRA, denitrification or nitrate reduction and the measurements of nitrite, ammonium, pH, and OD 600. Significant correlations, after FDR correction, are indicated by bold borders.

For each culture for which we have community composition data from 16S rDNA amplicons, we can sum the relative abundance of all strains possessing the genetic potential for nitrate reduction, DNRA or denitrification to estimate the total genetic potential for each of these nitrate respiratory traits. We observe that the genetic potential for DNRA is positively correlated with ammonium production and negatively correlated with nitrite production (Figure 4C, Figure 4D). This is largely driven by changes in the relative abundance of the dominant nitrate ammonifying *Escherichia* relative to the nitrite accumulating *Klebsiella*, but the *Citrobacter* and *Sulfurospillum* strains also contribute to DNRA genetic potential (Table S2).

### Specific carbon sources influence nitrate respiration end-products by selectively enriching for microbial sub-populations with distinct functional traits

To understand the basis for the selective enrichment of specific strains on different carbon sources, we profiled the carbon utilization capability of several isolates derived from the Jewel Lake enrichment culture. The highest fold-enrichment of strains relative to the dominant *Escherichia sp.* JL983 is for carbon sources that the enriched strains are uniquely capable of utilizing (Figure 5A). For example, *Klebsiella sp.* JL973 is the only isolated strain able to grow on L-sorbose or D-cellobiose and this strain is therefore highly enriched on these substrates. Similarly, the *Pseudomonas sp.* JL972 is the only isolated strain able to grow on citrate or formate and is likewise enriched on these substrates (Figure 5A).

**Figure 5.**
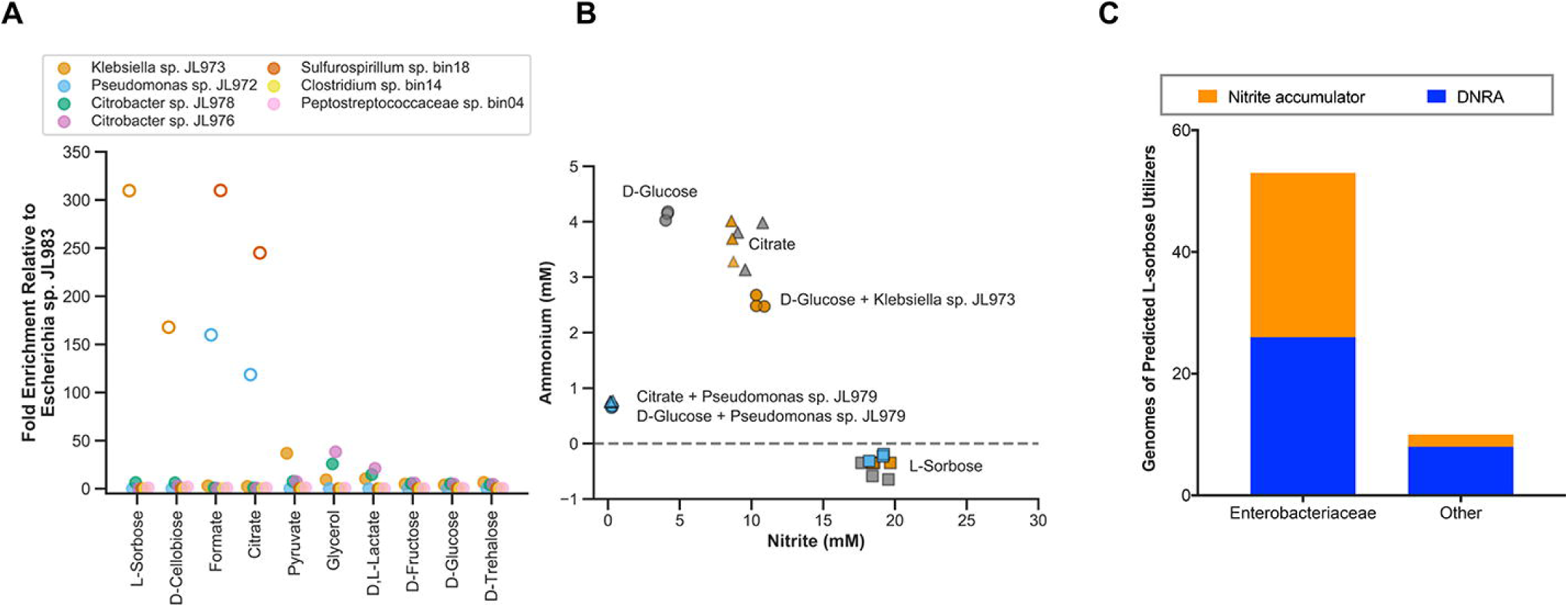
Selective carbon sources enrich for strains with distinct functional traits. **A.** Fold enrichment of strains relative to *Escherichia* in the Jewel Lake enrichment. Open symbols are strains that use carbon sources the *Escherichia* cannot utilize. **B.** Nitrite and ammonium concentrations of the Jewel Lake enrichment alone (dark grey symbols) or bioaugmented with *Klebsiella* (orange symbols) or *Pseudomonas* (blue symbols) with D-glucose, citrate or L-sorbose as the sole carbon source. **C.** For genomes that encode the L-sorbose utilization genes (*sorABE*), we show how often they are expected to be nitrite accumulators or nitrate ammonifiers (DNRA) based on the presence or absence of respiratory nitrate reductase genes (*napA, narG*) and respiratory nitrite reductase genes (*nirS, nirK, nrfA*). Because L-sorbose utilization is best studied in Enterobacteriaceae, we show results separately for this group than for other prokaryotes (Other). We found no predicted dentrifiers that have the *sorABE* genes. Data is from 27,941 prokaryotic genomes on Annotree (Materials and Methods).

To confirm the functional traits of the strains in the JL enrichment and demonstrate the selectivity of different carbon sources in influencing community functional outcomes, we bioaugmented the Jewel Lake enrichment with the *Klebsiella sp.* JL973 or *Pseudomonas sp.* JL972 strains at a 1:1 ratio of isolate to enrichment (Figure 5B). As expected, bioaugmentation with the *Klebsiella* strain shifts end-products towards more nitrite production and less ammonium production on D-glucose. There is no influence of *Klebsiella* sp. JL973 bioaugmentation on nitrate reduction end-products in L-sorbose cultures because *Klebsiella* sp. JL973 is already dominant on this carbon source and nitrate is stoichiometrically converted to nitrite.

There was no influence of *Klebsiella sp.* JL973 bioaugmentation in citrate cultures, because the *Klebsiella* strain does not utilize citrate. In contrast, *Pseudomonas* bioaugmentation in D-glucose and citrate cultures shifts end-products towards lower nitrite and ammonium production. This is consistent with the *Pseudomonas* strain’s capacity for complete denitrification and utilization of these carbon sources. While difficult to predict based on gene content, citrate utilization is heterogeneously distributed within the Enterobacteriaceae(*13, 14*), and anaerobic oxidation of citrate to formate by *Enterobacterial* isolates is generally not coupled to growth(*51*). In contrast, *Pseudomonas* and *Sulfurospirillum* are generally capable of utilizing citrate anaerobically coupled to nitrate reduction(*52*).

We also looked for the presence of L-sorbose utilization genes (*sorABE*) in the genomes of organisms predicted to be denitrifiers, nitrate ammonifiers or nitrite accumulators based on the presence or absence of nitrate reductase (*narG, napA*) and nitrite reductase (*nirS, nirK, nrfA*) genes (Figure 5C). The function of the *sorABE* genes has mostly been studied in the Enterobacteriaceae (*49*), and there may be other catabolic pathways for L-sorbose. However, a roughly equivalent number of predicted nitrate ammonifying (DNRA) and nitrite accumulating Enterobacteriaceae have *sorABE.* Thus, the enrichment of Enterobacterial nitrite accumulators, such as *Klebsiella* sp. JL973, may be a likely outcome after L-sorbose amendment in terrestrial environments. Taken together, our results demonstrate a selective influence of carbon sources in altering nitrate reduction end-products in the Jewel Lake enrichment by enriching for strains with specific carbon catabolic and nitrate respiratory traits.

## Discussion

A predictive understanding of how environmental perturbations influence the microbially-mediated nitrogen cycle has major implications for sustainable agriculture, wastewater treatment, and toxin remediation(*1, 2*). Previous work has demonstrated linkages between changes in carbon composition and microbial community composition(*21*-*24*), carbon composition and respiratory end-products(*5, 10, 53*), or community composition and respiratory end-products(*6*). However, few studies to date have examined the dynamics of nitrate respiring microbial communities using metagenomic sequencing (*6*), and rarely are the dynamics of genes, strains and respiratory traits systematically linked in a high-throughput format as we have in this study. A high-resolution understanding of how specific substrates impact the interactions and respiratory potential of specific microorganisms remains elusive, but will ultimately be required to model and predict the behavior of microbial communities and their ecosystem functions.

To overcome these challenges, in this study we applied a high-throughput approach to link diverse carbon sources to functional activity, community composition and genetic potential in nitrate-respiring microbial enrichment cultures. We found that specific carbon sources favor different end-products of nitrate respiration across three different microbial communities from geographically and geochemically distinct environments (Figure 2). To understand the mechanistic basis for these findings we sequenced the genomes of all dominant strains in an enrichment culture from aquatic sediment and identified correlations between dominant strains with different respiratory traits and the end-products of nitrate reduction (Figure 4). For example, the nitrate ammonifier, *Escherichia sp.* JL983 dominates on many sugars, but the nitrite accumulator, *Klebsiella sp.* JL973, is specifically enriched on L-sorbose and D-cellobiose and correlated with high nitrite concentrations and low ammonium concentrations. D,L-lactate, pyruvate and glycerol enrich for nitrate ammonifying *Citrobacter sp.* JL976 and *Citrobacter sp.* JL978. On citrate or formate, denitrifying *Pseudomonas sp.* JL972 are enriched, though not to the same extent as the nitrate ammonifying *Sulfurospirillum sp.* bin18 (Figure 3).

Carbon concentration is widely recognized as an important control on the competition between DNRA and denitrification(*3, 6, 7, 11*), but our results add important nuance to this paradigm. In our experiments, all cultures were amended with high concentrations of carbon relative to nitrate, and while nitrate ammonifiers were enriched in many cultures with correspondingly high ammonium production, nitrite accumulators or denitrifiers were enriched on other carbon sources with lower ammonium production and higher nitrite accumulation. We conclude that, while the thermodynamic advantage of DNRA is important, high local concentrations of a single carbon source can enrich for non-DNRA microorganisms until low-abundance nitrate ammonifiers capable of using that carbon source can grow to an extent where they can exert a dominant influence on the end-products of nitrate respiration. Therefore, accurate prediction of the end-products of nitrate respiration requires knowing the carbon concentration, carbon composition as well as the relative abundances, carbon catabolic traits and nitrate respiratory traits of each member of the microbial community.

We postulate that given the uneven distribution of catabolic and respiratory pathways in microbial genomes, different carbon sources will selectively favor different respiratory end-products by enriching for microbial sub-populations with different carbon catabolic traits. It is likely that selective carbon source amendments will be more effective in shifting respiratory end-products in less complex microbial communities with less dispersal, such as our enrichments or industrial reactors, versus open environments like aquifers, agricultural soils or lake sediment. However, in any environment, specific carbon source amendments will, at least transiently, enrich for distinct sub-populations with distinct respiratory traits. Thus, we anticipate that high-throughput approaches may help identify prebiotic amendments that can influence nitrate respiratory end-products in industrial ecosystems, for example, to stimulate denitrification in wastewater treatment facilities or to stimulate DNRA in agricultural soils.

More broadly, correctly predicting the influence of diverse selective pressures on complex microbial communities with heterogeneous gene content and traits will require better functional annotations for genes and strains. While genome-resolved metagenomics provides high-resolution snapshots of microbial communities, high-throughput laboratory simulations are essential to understand how changing conditions influence community dynamics. There is much to be gained by combining these two approaches as we have in this study. By studying the dynamics and function of genomically-characterized, low-complexity microbial communities in high-throughput, we anticipate rapid advances in mechanistic ecology that will improve our ability to accurately predict the influence of complex, variable environmental parameters on microbially-mediated processes.

## Acknowledgements

This work was funded by ENIGMA, a Scientific Focus Area Program supported by the U. S. Department of Energy, Office of Science, Office of Biological and Environmental Research, Genomics: GTL Foundational Science through contract DE-AC02-05CH11231 between Lawrence Berkeley National Laboratory and the U. S. Department of Energy. The authors would like to thank members of the ENIGMA Scientific Focus Area as well as John D. Coates (UC. Berkeley), Tyler C. Barnum (UC Berkeley) and David C. Vuono (Colorado School of Mines) for helpful feedback and suggestions.

## Data Availability

DNA sequencing data are available under BioProject Accession PRJNA576510.

## Supplemental materials

**Figure S1. A.** Nitrite concentrations and donor electron equivalents for the Jewel Lake enrichment recovered on 10 different carbon sources in triplicate. **B.** Nitrite concentrations and donor electron equivalents for the Jewel Lake enrichment recovered on 10 different carbon sources in triplicate. **C.** Ammonium concentrations and pH for the Jewel Lake enrichment recovered on 10 different carbon sources in triplicate. **D.** Ammonium concentrations and pH for the Jewel Lake enrichment recovered on 10 different carbon sources in triplicate. Points are colored based on which dominant strain is most highly selectively enriched in each condition.

**Figure S2.** Ammonium (A-E) or nitrite concentrations in the Jewel Lake enrichment recovered on different carbon sources plotted against relative abundances of *Escherichia* and *Klebsiella* (A), *Citrobacter* (B), *Pseudomonas* and *Sulfurospirillum* (C), or *Clostridium* and *Peptostreptococcaceae* (D) strains.

## Dataset S1

**Table S1.** Carbon source influence on the end-products of nitrate reduction in enrichment cultures.

**Table S2.** Taxonomy and genetic functional potential of strains in Jewel Lake enrichment culture

**Table S3.** Functional activity and community composition of Jewel Lake enrichment recovered on various carbon sources.

**Table S4.** MetaBat2 and CheckM results and nitrogen cycling genes for metagenome assembled genomes from Jewel Lake enrichment

