## Supplementary figures and images for "Selective carbon sources influence the end-products of microbial nitrate respiration"

### Figure S1

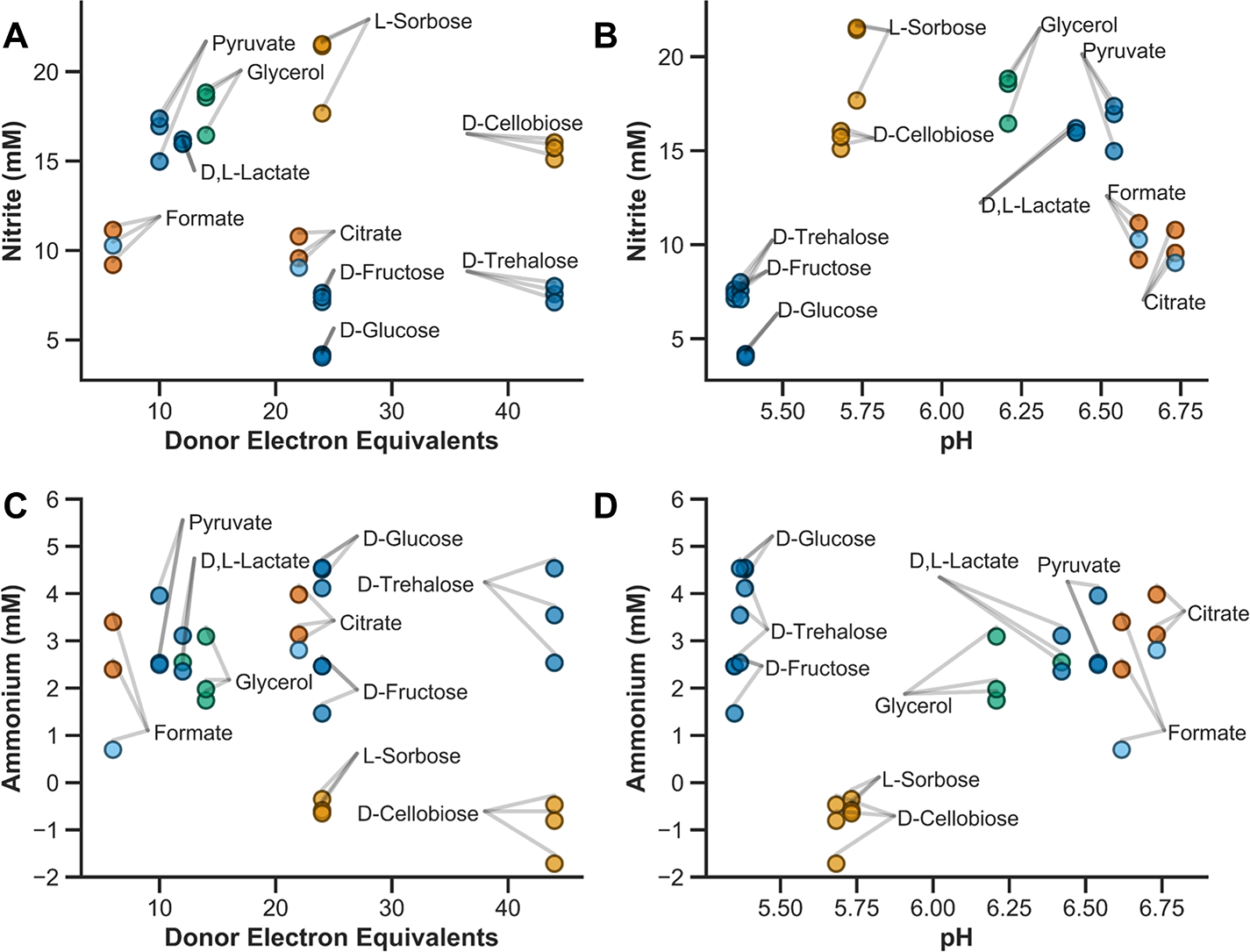

### Figure S2

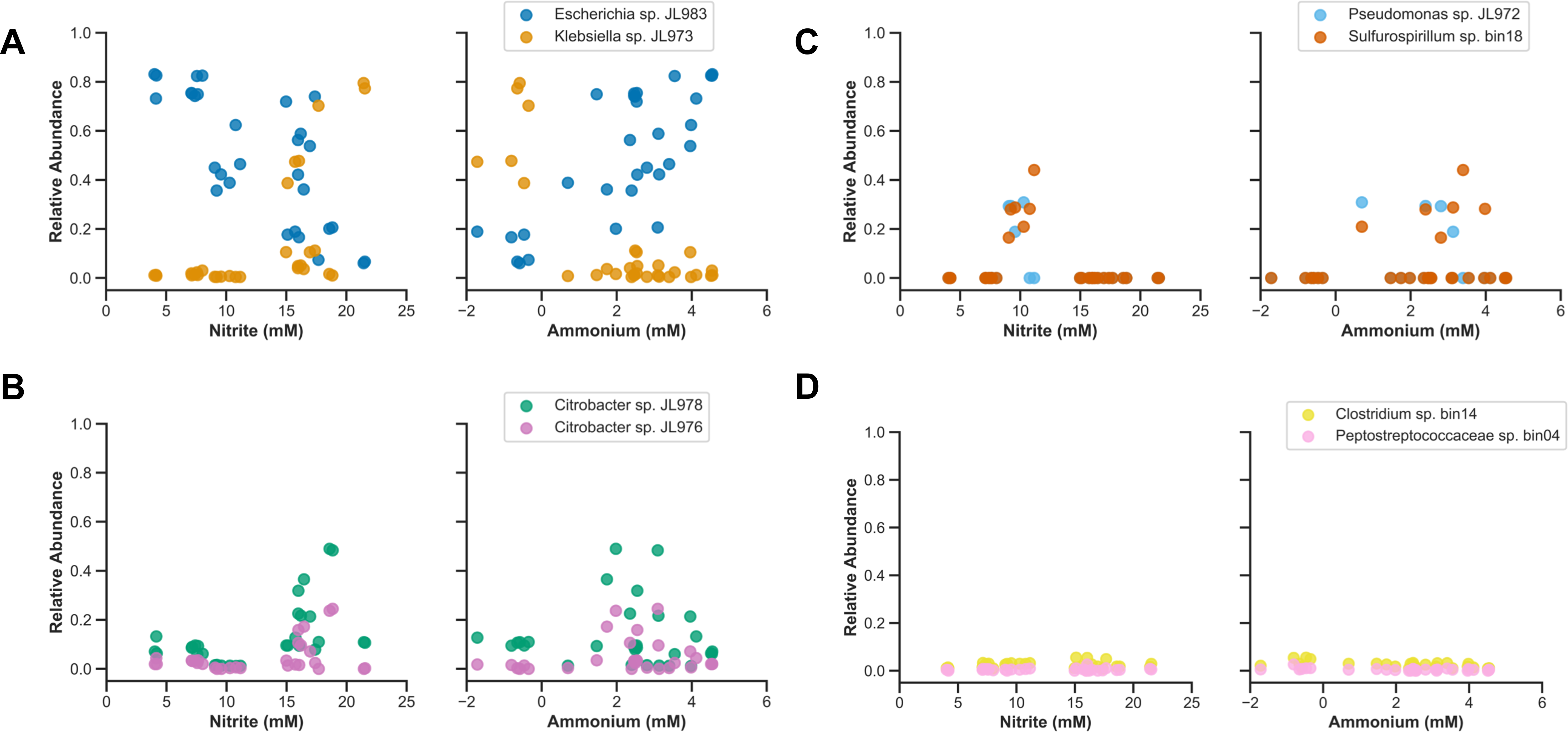
